# Functional characterization of retinal ganglion cells using tailored nonlinear modeling

**DOI:** 10.1101/421396

**Authors:** Qing Shi, Pranjal Gupta, Alexandra Boukhvalova, Joshua H. Singer, Daniel A. Butts

**Affiliations:** Department of Biology, University of Maryland, College Park, MD, United States; Program in Neuroscience and Cognitive Science, University of Maryland, College Park, MD, United States

## Abstract

There are 20-50 functionally- and anatomically-distinct ganglion cell types in the mammalian retina; each type encodes a unique feature of the visual world and conveys it via action potentials to the brain. Individual ganglion cells receive input from unique presynaptic retinal circuits, and the characteristic patterns of light-evoked action potentials in each ganglion cell type therefore reflect computations encoded in synaptic input and in postsynaptic signal integration and spike generation. Unfortunately, there is a dearth of tools for characterizing retinal ganglion cell computation. Therefore, we developed a statistical model, the separable Nonlinear Input Model, capable of characterizing the large array of distinct computations reflected in retinal ganglion cell spiking. We recorded ganglion cell responses to a correlated noise (“cloud”) stimulus designed to accentuate the important features of retinal processing in an *in vitro* preparation of mouse retina and found that this model accurately predicted ganglion cell responses at high spatiotemporal resolution. It identified multiple receptive fields (RFs) reflecting the main excitatory and suppressive components of the response of each neuron. Most significantly, our model succeeds where others fail, accurately identifying ON-OFF cells and segregating their distinct ON and OFF selectivity and demonstrating the presence of different types of suppressive receptive fields. In total, our computational approach offers rich description of ganglion cell computation and sets a foundation for relating retinal computation to retinal circuitry.

## INTRODUCTION

The first stages of sensory processing encode sensory input in a format that allows higher brain areas to extract information essential to guide behavior. In the visual system, tens of different types of ganglion cells— the retinal output neurons — serve as “feature detectors” that each encode specific components of the visual world and convey information to the brain (Lettvin et al., 1959; Barlow and Levick, 1965; Olveczky et al., 2003; Münch et al., 2009; Dhande and Huberman, 2014; Dhande et al., 2015). Feature detection is accomplished by distributing the output of rod and cone photoreceptors to a series of parallel neural circuits, each of which originates with one of ∼15 types of bipolar cell and provides input to multiple types of ganglion cell (Masland, 2012; Demb and Singer, 2015). Thus, the responses of different classes of ganglion cells thus reflect computations performed by the various presynaptic, inner-retinal, parallel circuits (Euler et al., 2014). Although the anatomy and retinal circuitry supporting the diversity of ganglion cell classes has been studied in molecular (Helmstaedter et al., 2013; Sümbül et al., 2014), physiological (Baden et al., 2016), and statistical and computational detail (Carcieri et al., 2003; Segev et al., 2006; Seung and Sümbül, 2014), the computational methodology necessary to represent the diversity of retinal computations reflected in ganglion cell outputs is relatively anemic.

Being able to describe the “diversity of computation” requires sufficiently complex mathematical descriptions of how each ganglion cell processes visual stimuli. The most commonly used mathematical model of the retina is the linear-nonlinear (LN) model, which describes the transformation from a visual stimulus to action potentials as a two-step process: encompassing first a linear filter that emphasizes particular spatial and temporal features in the stimulus; and second, a nonlinearity that accounts for action potential generation (Chichilnisky, 2001). The LN model is appealing in its simplicity and yields an easily interpretable spatiotemporal filter (also called the linear receptive field, or RF), which typically has a center-surround structure and either ON or OFF selectivity in time (i.e. its polarity indicates whether neurons respond to increases or decreases in luminance). The second stage, “spiking nonlinearity”, describes how sensitively neuron firing rates depend on the stimulus (Kim and Rieke, 2001; Chichilnisky and Kalmar, 2002; Zaghloul et al., 2005).

Although the LN model can distinguish ON from OFF cells based in its RF, this single linear filter condenses all of retinal processing into a single step, which masks the diversity of processing channels presynaptic to ganglion cells (Asari and Meister, 2012). This is a particular problem in the mouse retina, where ON-OFF cells (i.e., cells that respond both to increments and decrements in light intensity) are the most common cell type (Zhang et al., 2012). ON-OFF cells are described as having least two receptive fields (one ON and one OFF), and a single linear receptive field therefore averages ON and OFF filters together (Kim and Rieke, 2001; Chichilnisky and Kalmar, 2002; Cantrell et al., 2010; McFarland et al., 2013). Likewise, direct inhibitory inputs onto ganglion cells (which is nearly ubiquitous) would also not be able to be distinguished using linear RFs (Butts et al., 2011), and furthermore would be unable to distinguish OFF inhibition with ON excitation.

Thus, more sophisticated mathematical models are required to capture the diverse array of retinal computations. Such models are necessarily nonlinear, and they involve the characterization of multiple receptive fields corresponding to diverse excitatory and suppressive features (Schwartz et al., 2012; McFarland et al., 2013; Freeman et al., 2015; Liu et al., 2017; Maheswaranathan et al., 2018). Nonlinear models in general are much more difficult to fit to data because they require many more parameters (e.g., multiple RFs) and require a nonlinear description that can capture a sufficient range of retinal computation. Here, we use a “space-time separable” form of the Nonlinear Input Model (NIM) (McFarland et al., 2013) to fit multiple excitatory and suppressive RFs by maximum-likelihood modeling. Space-time separability refers to representing spatiotemporal RFs as combinations of separate spatial and temporal filters; this permits much more efficient parameter estimation (McFarland et al., 2013; Park and Pillow, 2013; Thorson et al., 2015; Maheswaranathan et al., 2018). Furthermore, because this approach does not require uncorrelated “white noise” stimuli, we can characterize ganglion cells experimentally using tailored “cloud” stimuli, which have spatial features on multiple scales.

The NIM described detailed spatiotemporal RF maps in ON, OFF, and (uniquely) ON-OFF ganglion cells in the mammalian retina, and it revealed suppressive RF components not observed with standard LN analyses. Such detail provides a much fuller picture of the computations represented in ganglion cell outputs and provides the means to understand their functional diversity.

## RESULTS

We recorded retinal ganglion cells from *in vitro* mouse retina using a multielectrode array. We first consider recordings made during the presentation of spatiotemporal white noise in order to perform standard receptive field (RF) analyses. The RF of ganglion cells is usually defined to be the optimal linear filter **k** that best predicts the spike response of the neuron (Dayan and Abbott, 2001), and is the basis of the Linear-Nonlinear (LN) model, which generates a firing rate prediction of the response *r*(*t*) given the stimulus **s**(*t*) as follows (Fig. 1A):

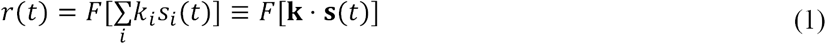

where *i* indexes the components of **s**(*t*) and the corresponding filter **k** that are relevant to the response at time *t* (including spatial locations and time lags into the past), and *F*[.] is a spiking nonlinearity that maps the output of the filter to a firing rate. Note we are using boldface to represent vectors, i.e., **k**=[*k*_1_ *k*_2_ *k*_3_ …].

**Figure 1:**
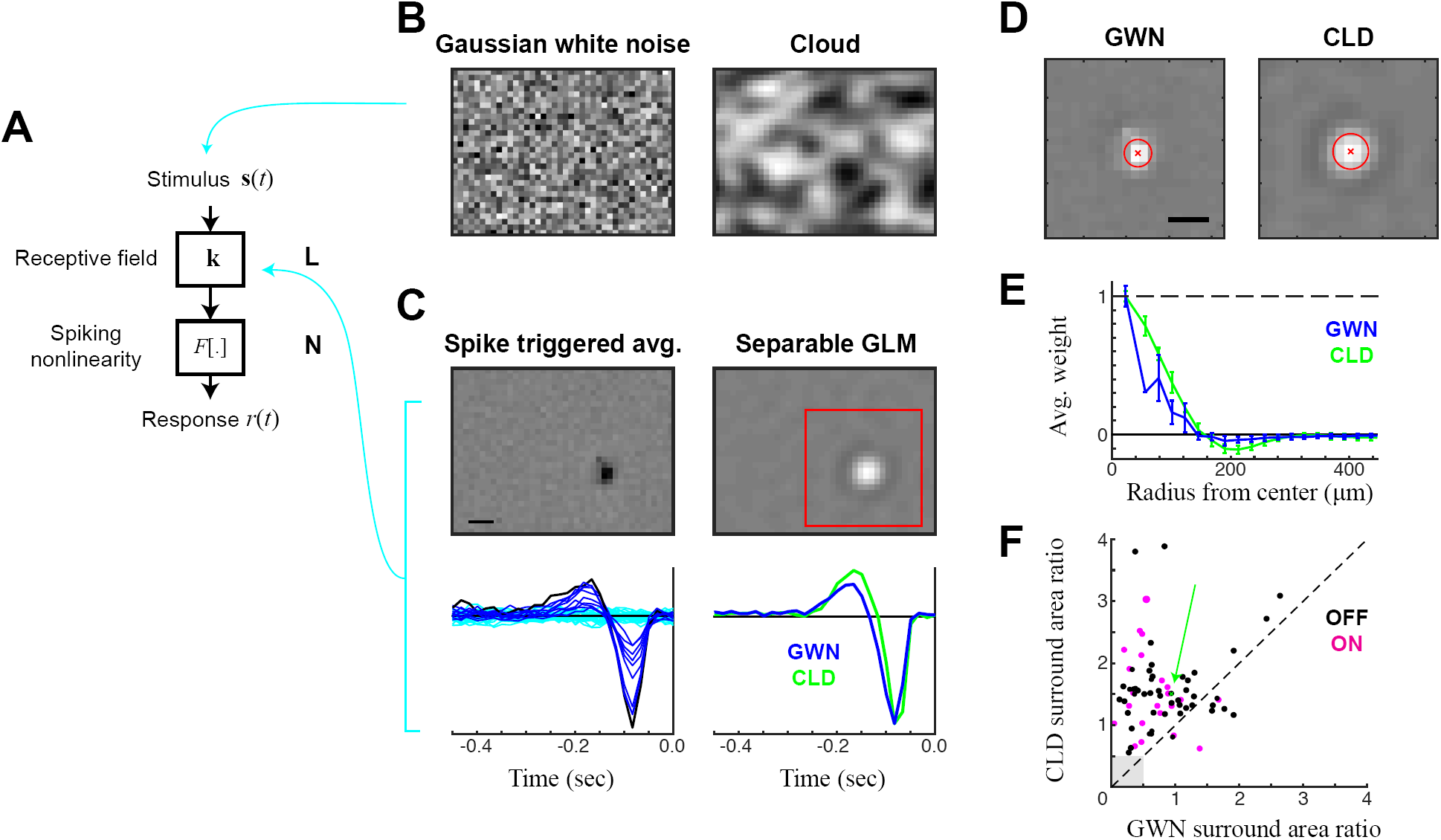
Estimation of spatiotemporal RFs using separable GLM and cloud stimuli. **A.** Schematic for the Linear-Nonlinear (LN) model of ganglion cells: consisting of a linear spatiotemporal filter **k**, followed by a spiking nonlinearity *F*[.]. **B.** Example frames from the two stimulus classes used for model estimation: gaussian white noise (GWN, *left*) and cloud stimuli (CLD, *right*). (**C-E**) Analysis of an example OFF cell presented with both GWN and CLD stimuli. **C.** LN model filters, fit using spike-triggered averaging (STA) applied to GWN data (*left*), compared with a space-time separable generalized linear model (GLM) fit to the cloud stimulus (*right*). The STA yields a temporal function for every pixel (*bottom left*), from which the spatial map at the best latency can be extracted (*top left*). By comparison, the separable GLM extracts a single temporal function (*bottom right*), which is multiplicatively combined with a single spatial function (*top right*). This reveals a much less noisy spatial estimate, from which an opposite-sign surround is visible. The temporal kernel is also shown for the separable GLM fit to the GWN stimulus, revealing a slightly longer-latency response, but otherwise very similar temporal tuning. **D.** Spatial maps derived from a separable GLM applied to the GWN (*left*) and CLD (*right*) stimuli, revealing the ability of the CLD stimuli to drive clearer spatial information at larger scales: ‘x’ shows the estimated center, and one standard deviation shown by the red circle. **E.** To measure the presence of the spatial surround, the average pixel intensity (and standard deviation) at each radius is plotted versus the radius. This also demonstrates the ability of the CLD stimulus (green) to more robustly recover the surround. **F.** The surround-area-to-center-area ratio for CLD vs. GWN stimuli, shown for each ON and OFF cell where both conditions were recorded (*N*=81, where 7 cells with negligible surround areas (gray square) not shown). Scale bars are 200 µm.

When the stimulus is uncorrelated across its dimensions, then the optimal linear filter can be simply estimated using the spike triggered average (STA) (Chichilnisky, 2001; Dayan and Abbott, 2001), defined as the average stimulus that evoked a spike. The spiking nonlinearity also can be estimated easily using the histogram (Chichilnisky, 2001; Dayan and Abbott, 2001). For this reason, the LN model estimated in the context of Gaussian white noise (GWN; Fig 1B, *left*) is the predominant method used to characterize ganglion cells (Chichilnisky, 2001).

In using a white noise stimulus, however, consideration of the spatial extent and temporal duration of the pixels that compose the stimulus is critical (Meister et al., 1994; Pamplona et al., 2015). Spatial and temporal parameters often are chosen empirically either to drive strong ganglion cell responses or to yield RFs with good appearances, or both. A good RF for ganglion cells usually has a relatively smooth center covered by a few pixels (Fig. 1C, *left*); note, though, that in many publications the RFs pictured are interpolated into a higher resolution image and smoothed. While, in principle, smoother RFs can be obtained with finer spatial resolution using smaller pixels, having a large number of uncorrelated pixels tiling any one feature (e.g., theRF center) will typically not drive robust neuron responses, because the summed luminance over that feature will not deviate much from gray (mean) luminance. In the context of MEA recording, where RFs of many sizes might be sampled, the stimulus is usually adjusted to the most prevalent RF sizes, and therefore can be maladjusted for others (Pamplona et al., 2015).

Furthermore, even an optimal choice of pixel size for a given RF center can bias RF estimation to features of that size. RF features that have larger scales, such as the RF surround, will not be driven effectively by GWN stimuli optimized for the center: again, this occurs because the uncorrelated light and dark pixels will average together to gray and not drive the larger features of the RF. As a result, features such as the RF surround will contribute minimally to the response to white noise stimulation and often not be visible in the spike-triggered average (Fig. 1C, *left*), which is a common issue with white noise characterizations throughout previous work.

### Likelihood based estimation of RFs using cloud stimuli

We presented a “cloud” stimulus to the same neurons characterized using GWN. This cloud stimulus was designed to have the same pixel size and duration as the GWN stimulus, but it introduced spatial correlations within each frame; these were generated by applying a spatial low-pass filter to a GWN stimulus (see Methods). Such correlations implicitly lead to areas of dark and bright areas at a range of spatial scales (Fig. 1B, *right*) while maintaining the same high spatial resolution as GWN.

The introduction of such correlations confounds traditional spike-triggered characterizations because the presence of correlations requires an additional (and noisy) step in computing the optimal linear filter: deconvolution of stimulus correlations from the STA (Dayan and Abbott, 2001; Theunissen et al., 2001). This problem can be avoided by direct estimation of the linear filer by maximum *a posteriori* estimation, such as in the framework of the Generalized Linear Model (GLM) (Paninski, 2004; Truccolo et al., 2005). Such optimization is easily performed in principle (given standard computers and software), but it can introduce problems of overfitting due to the large number of parameters in the spatiotemporal filter that must be estimated simultaneously. For example, an appropriate spatial and temporal resolution for our experiments (21×21 spatial grid and 40 time lags at 16 ms resolution) results in thousands of parameters (17,640 in this case) that must be fit simultaneously. Application of smoothness regularization can mitigate overfitting, but its ability to suppress the noise in the RF without over-smoothing is limited, given typical amounts of data. Thus, in the context of relatively high-resolution spatiotemporal estimation problems, the GLM often provides little (if any) advantage over the STA in the context of a GWN stimulus; STA estimation is not affected by the number of parameters because each element is an independently performed average.

This major limitation of GLM estimation of ganglion cell RFs can be overcome by low-rank approximation (Park and Pillow, 2013; Thorson et al., 2015; Maheswaranathan et al., 2018), which involves representing the spatiotemporal RF as a combination of separately estimated spatial and temporal RFs. Therefore, we adapted GLM estimation techniques to estimate space-time separable filters by representing the spatiotemporal filter as a spatial filter multiplied by a temporal filter, i.e., *k*(*x,y*,τ) = *k*_*sp*_(*x,y*)×*k*_t_(τ). This separable form is fit through block-coordinate descent: after initializing with the [correlated] STA, the spatial filter is held fixed while the temporal kernel is estimated (40 parameters), and then the temporal kernel is held fixed while estimating the spatial kernel (441 parameters). Alternating continues until the fit converges (see Methods), and corresponding spatial and temporal regularization is applied to each separately. Such an approach yields clean estimates of both (Fig. 1C, *right*): with much smaller amount of noise than the STA, and the appearance of additional details of the RF such as an opposite-polarity surround.

The separable GLM can be applied to both GWN and cloud data, yielding nearly identical temporal kernels (Fig. 1C). RFs measured from the cloud stimulus context, however, yield much more consistent features (Fig. 1D). Note that both RFs are optimized in their respective contexts: differences in spatial structure represent the spatial patterns that best predicted responses using an LN model. Although the neuron shown fired roughly the same number of spikes in each context (58,219 in GWN and 61,358 in CLD), RF elements of the cloud stimulus drive the ganglion cell much more efficiently, leading to more robust features in the estimated RF (Fig. 1D) including a smaller latency to spike (Fig. 1C, *bottom right*). This is most evident in the appearance of the RF surround, which we measured by fitting the center to a circular Gaussian (Fig. 1D, red) and then measuring the average value of the RF at concentric distances to yield a precise measurement of its strength (Fig. 1E). While there was a large diversity in surround strength for the LN models measured across neurons, the cloud stimulus condition yielded stronger surrounds in all but a handful of cases (Fig. 1F).

### Estimation of ON-OFF receptive fields

A large fraction of ganglion cells in mouse are ON-OFF, meaning they are excited by both increments and decrements in luminance (Fig. 2A). Such selectivity implicitly defies characterization with linear RFs: linear stimulus processing implicitly averages the selectivity of different RF components into a single RF, and overlapping ON and OFF components cancel, i.e., (**k**_*on*_ · **s**) + (**k**_*off*_ · **s**) = (**k**_*on*_+ **k**_*off*_) · **s**. For two RF components to contribute separately to excitation, ON and OFF selectivity must be represented in an additional nonlinear stage of processing (Fig. 2A):

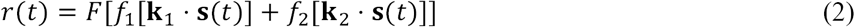

**Figure 2:**
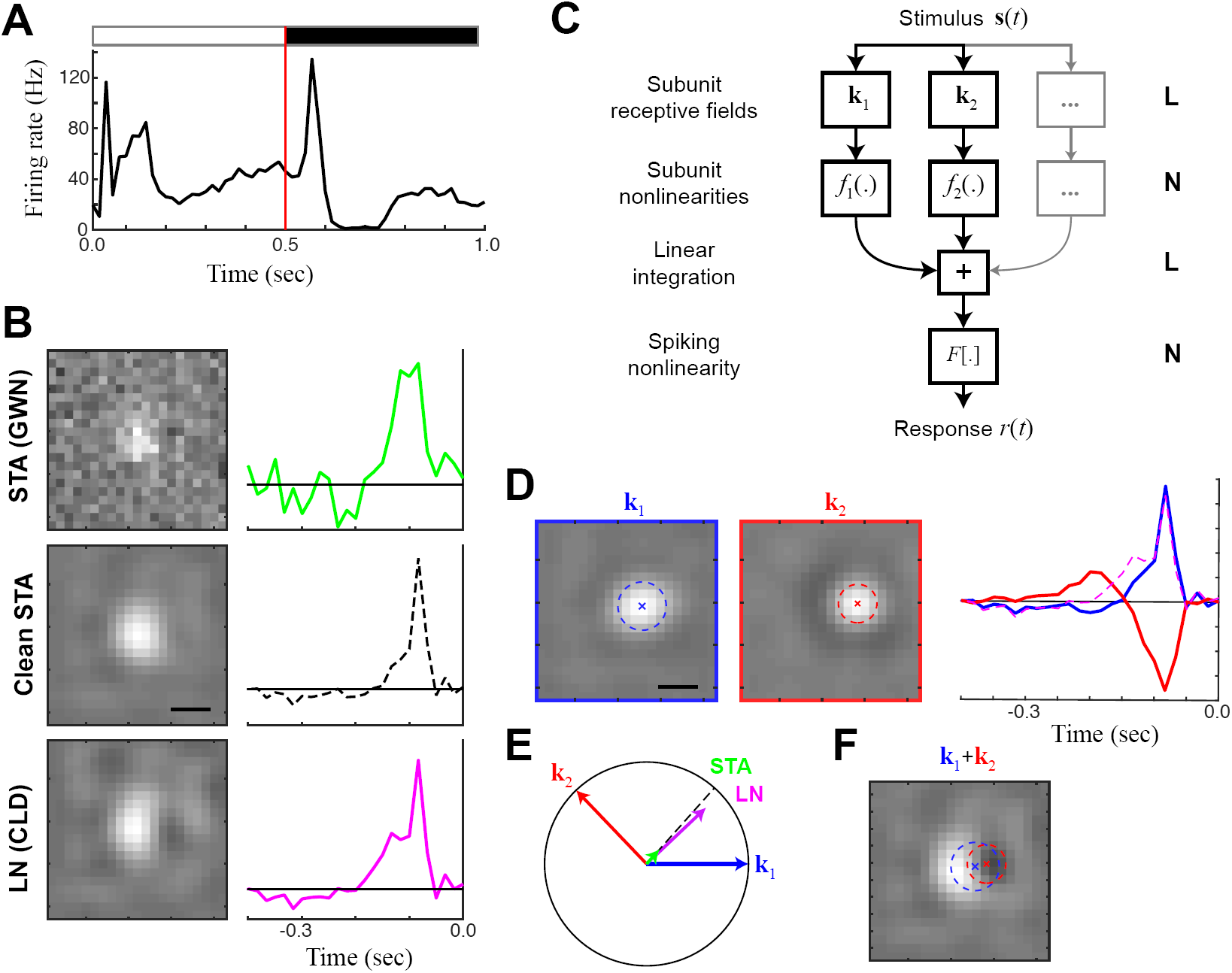
Receptive field mapping of ON-OFF cells. **A.** The PSTH of an example ON-OFF cell in response to a full-field flashed stimulus alternating between black and white at 2 Hz. **B.** The LN model (estimated using the STA and separable GLM) are limited to compute a single receptive field for the cell. As with Figure 1, the separable GLM (bottom) reveals the most spatial detail relative to the STA in GWN (*top*), although can be ‘cleaned’ (*middle*, see E) to reveal a similar receptive field as the GLM. **C.** The structure of the NIM model, which is an LN-LN cascade that can identify several features (each with its own receptive field) that nonlinearly combine to result in the spike output. **D.** The separable NIM (sNIM) finds two excitatory receptive fields for the ON-OFF cell. By convention, the spatial map (*left*) is always positive, and the temporal kernels (*right*) demonstrate that one is ON and the other OFF. **E.** The ON and OFF spatiotemporal filters of the sNIM can be considered vectors in high-dimensional stimulus space, and define a plane, with the circle showing unit-length in the plane that denotes whether a given filter fully projects into the plane. The LN model filter (magenta, from B) projects largely into this plane at the average position between the two sNIM vectors. Because of noise, the STA (green, also from B) largely projects outside of the plane, but its projection into the plane matches the LN filter, and can be “cleaned” by normalizing this in-plane projection. **F.** As suggested by (E), simply averaging the ON and OFF NIM filters together produces a spatial map very similar to that of the LN model, with almost unnoticeable dark patch a result of the spatial offset of the OFF receptive field relative to the ON receptive field. Scale bars are 200 µm.

This model has the form of an LNLN cascade (Hochstein and Shapley, 1976; Korenberg et al., 1989; Butts et al., 2011) and has been formulated in a maximum likelihood context as the Nonlinear Input Model (NIM) (McFarland et al., 2013).

We adapted the NIM to incorporate separable spatiotemporal filters, and the resulting model, the *separable NIM* (sNIM) was fit to the same data as the separable GLM described above. To begin, we show an example cell that responds to both increments and decrements in luminance (i.e., ON and OFF stimuli) in a slowly modulated full-field stimulus (Fig. 2B). The selectivity demonstrated by the STA suggested that this cell is an ON cell because the calculation is limited to computing a single RF. The separable GLM, however, provided additional resolution and revealed a spatial asymmetry in the spatial RF (*right*), suggesting spatially offset and partially cancelling ON and OFF components. The sNIM fit to the same data demonstrated robust, circular ON and OFF responses, each with a discernible surround (Fig. 2C). The resulting two filters are represented as vectors that define a plane (Fig. 2E) in the much higher-dimensional “stimulus space”, which represents all possible types of feature selectivity. This plane defined by the detected ON and OFF filters largely contains the GLM filter, which in this case matched the sum of the two NIM filters (Fig. 2F). Although most of the power in the STA is dominated by noise, its projection into this plane is also in the same direction as the GLM (Fig. 2B, *middle*).

The sensitivity of the sNIM reveals ON and OFF components of the response in ganglion cells that otherwise would appear to be exclusively ON or OFF cells. ON-OFF cells are often characterized based on response to increments and decrements in full-field luminance (e.g., Fig. 2A). Measures of the peak responses to ON and OFF steps (Carcieri et al., 2003) yield a distribution of response types (Fig. 3A, blue), ranging from pure ON to pure OFF with a large number of intermediate values. Such measures, however, are poor indicators of true ON-OFF cells, as demonstrated by the histogram of values for neurons with clear ON and OFF excitatory components in the sNIM (red). For example, we show an example neuron having a negligible response to a full-field decrement in luminance (Fig. 3B) and a clear ON RF predicted by the GLM (Fig. 3C). This cell, however, is best described by an ON-OFF sNIM (Fig. 3D), and its ON-OFF selectivity can be demonstrated from analysis of responses to repeated presentations of a unique cloud stimulus (Fig. 3G).

**Figure 3:**
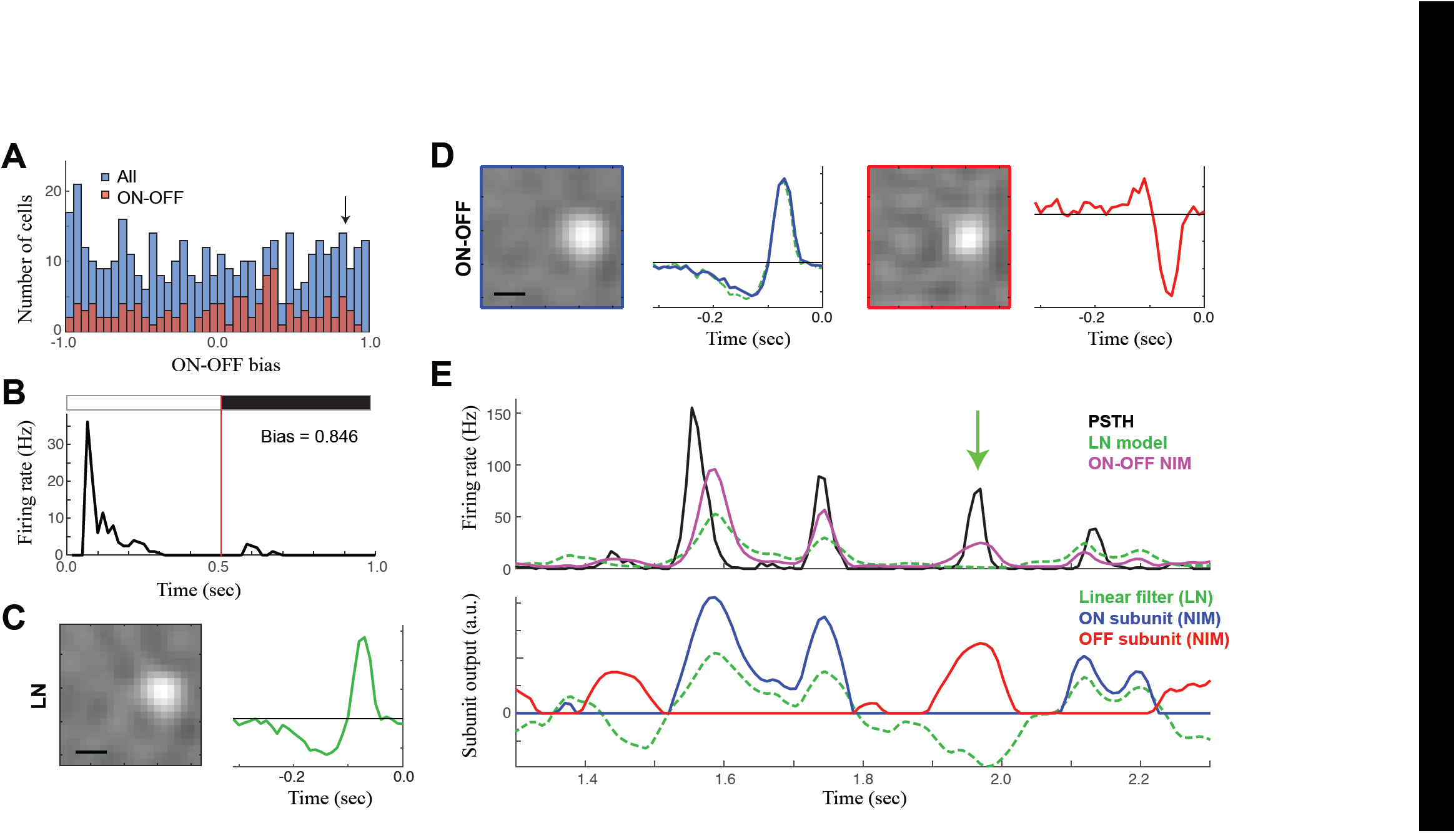
Hidden ON-OFF cells. **A.** ON-OFF cells are often distinguished using their response to full-field flashes (as in Fig. 2A), leading to a simple measure of ON-OFF bias that is simply the normalized difference in response magnitude between ON and OFF flashes. This leads to a continuum of values across all ganglion cells (blue, *N=XX*). However, using the sNIM to discriminate ON-OFF cells based on the presence of excitatory ON and OFF subunits, yields a similar continuum of values for ON-OFF bias (red). **B.** The PSTH in response to full-field flashes of an example ON-OFF cell identified by the sNIM, but has a high ON-OFF bias consistent with an ON cell. **C.** The same example cell’s linear receptive field is also consistent with an ON classification. **D.** However, the sNIM pulls and ON and OFF excitatory receptive field. The ON receptive field nearly matches that of the LN model (green). E. To demonstrate it is indeed an ON-OFF cell, the response to repeated presentations of the CLD stimulus is shown (*top*), relative to predictions of the ON-OFF sNIM (purple) and LN model (green). The underlying output of each subunit is shown *below*. The green arrow denotes an example of a response to an OFF stimulus, demonstrating it indeed has excitatory OFF responses. Scale bars are 200 µm.

We thus characterized neurons as being ON-OFF based on whether their best model (i.e., that with the highest cross-validated model performance) had two excitatory subunits with opposing polarity. We used the log-likelihood (cross-validated, *LL*_*x*_), which is ideal for long continuous trials because it does not require repeated stimuli to compute (as compared with the more common measure such as explainable variance *R*^2^), as a measure of model performance (Pillow et al., 2005; Butts et al., 2011; Cui et al., 2016a). The *LL*_*x*_-improvement for the ON-OFF model relative to the separable GLM was quite variable across labeled ON-OFF cells (Fig. 4A), with had a median improvement of 55% (*N*=141). There, however, were a large number of ON-OFF cells with small improvements (Fig. 4B). However, this was largely due to the distribution of ON-OFF bias (Fig. 4C), such neurons that were highly dominated by ON- or OFF-responses had little to gain by correctly predicting the response to opposite polarity, and thus large ON-OFF bias was associated with very little model improvement. While the PSTH-based analysis shown above (Fig. 3G) can demonstrate the clear presence of ON and OFF responses, such analysis was not possible with the most of the dataset, because it depended on having repeated stimuli for each neuron that happens to contain noise patterns match its ON and OFF RFs. As a result, model-based labeling cannot be definitive, but rather demonstrates the much more wide-spread presence of ON-OFF selectivity than what is traditionally reported through electrophysiological characterizations.

**Figure 4:**
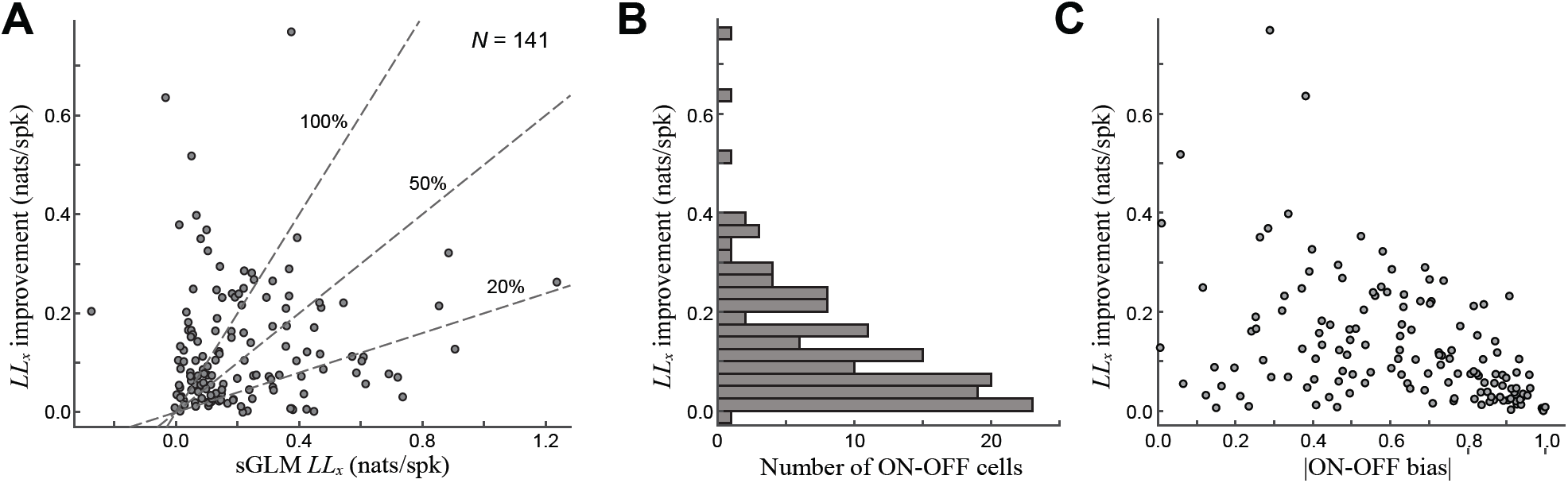
Cross-validated performance of ON-OFF sNIM. **A.** The cross-validated log-likelihood (*LL*_*x*_) improvement of the ON-OFF sNIM with two excitatory RFs (e.g., Figs. 2,3), relative to the *LL*_*x*_ of the separable GLM. **B.** A histogram of these *LL*_*x*_-improvements demonstrate most ON-OFF sNIM models contribute little to the model performance. **C.** However, this can be largely explained by the distribution of ON-OFF bias (absolute value shown on horizontal axis), such that ON-OFF cells that are dominated by ON or OFF component (i.e. not balanced), will tend to show smaller improvements.

### Detection of suppressive surrounds

Although ON-OFF cells are the most obvious examples of nonlinear processing in the retina, most ganglion cells exhibit some form of nonlinear processing in their responses, such as adaptation to stimulus contrast (Shapley and Victor, 1978; Brown and Masland, 2001; Kim and Rieke, 2001; Baccus and Meister, 2002) and the generation of temporal precision (Berry and Meister, 1998; Keat et al., 2001; Liu et al., 2001; Uzzell and Chichilnisky, 2004). In the context of the NIM model, such nonlinear properties might be explained through the addition of suppressive subunits (Bonin et al., 2006; Mante et al., 2008; Butts et al., 2011; 2016; Cui et al., 2016b), e.g., the same model structure that fit ON-OFF cells (Fig. 2C) with the second subunit’s output multiplied by −1 such that it can only suppress the firing rate. Indeed, a majority of OFF cells have a suppressive RFs identified by the sNIM (e.g., Fig. 5). These suppressive subunits typically have roughly the same spatial extent as the excitatory subunit (and LN model RF) (Fig. 5A, *left*), but with a delayed temporal kernel (Fig. 5A, *middle*). Such delayed suppression is consistent with similar models fit to cat retinal ganglion cells (Butts et al., 2016) and LGN cells (Butts et al., 2011; McFarland et al., 2013; Butts et al., 2016). Compared with the corresponding LN model, both the spatial and temporal filters of the NIM have broader dimensions; likely because the LN model’s RF the average of filters that largely cancel each other. The contribution of the suppressive term tends to make the response more transient due to the delay in the suppression (Butts et al., 2011).

**Figure 5:**
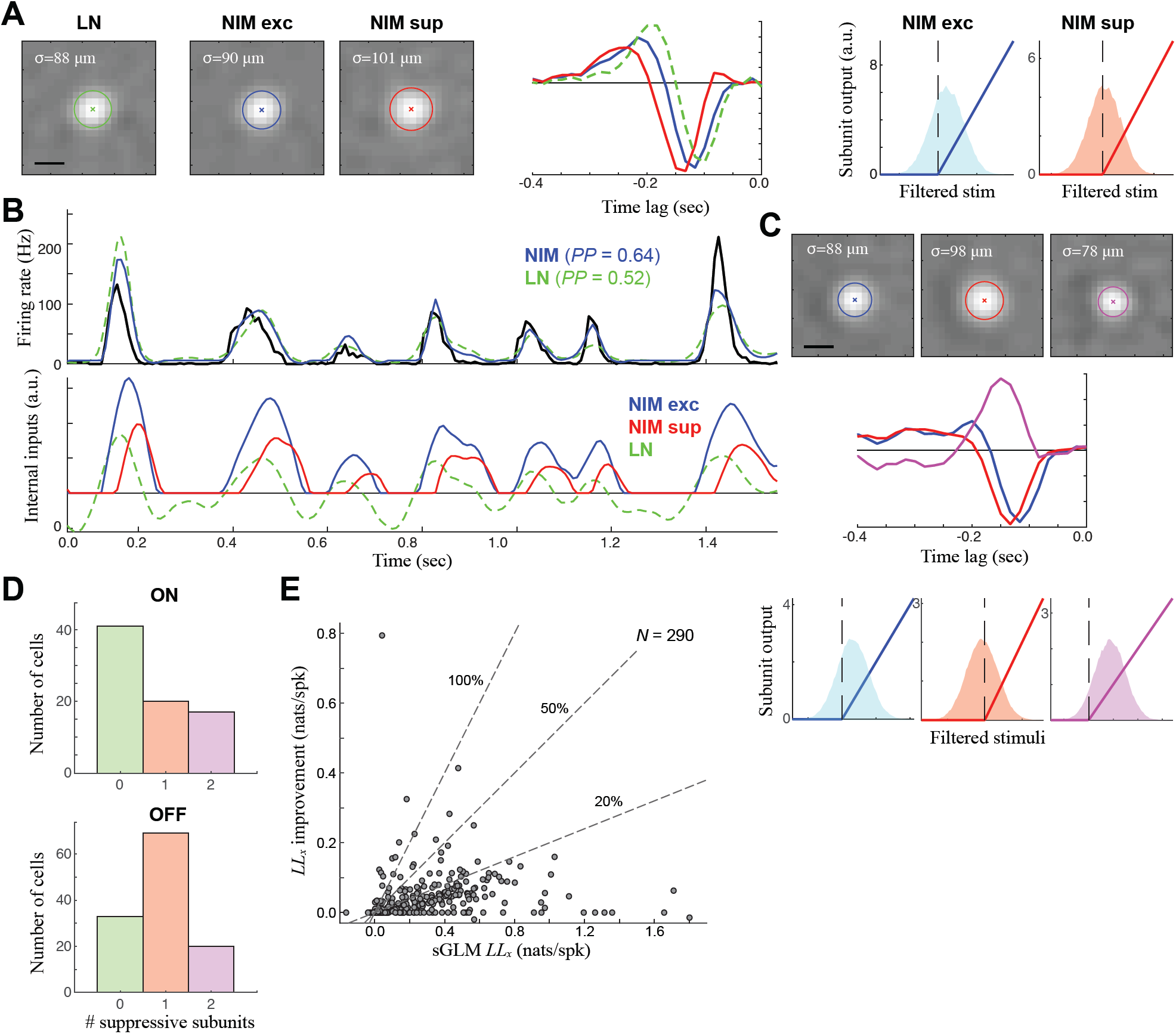
Detection of nonlinear suppression a majority ganglion cells. **A.** The sNIM model components for a typical OFF cell, which is best modeled by an excitatory (blue) and delayed suppressive (red) subunit. Both spatial components (*left*) are slightly larger than that predicted by the LN model, delayed in time relative to the LN model (*middle*), and strongly rectified (*right*). **B.** The resulting sNIM predicts the response better (*top*), although the underlying excitation and delayed suppression largely cancel (bottom) in order to offer only subtle differences between the LN model and sNIM. **C.** A different example OFF cell with two suppressive subunits: a delayed same-sign suppression (as with the example in A), as well as an opposite-sign ON suppression. **D.** The distribution of detected subunits for ON (left) and OFF cells (right), demonstrating that a majority of ON and OFF cells have underlying suppressive receptive fields in addition to standard excitatory subunits that largely confer their tuning. **E.** The cross-validated log-likelihood (*LL*_*x*_) improvement of adding suppression across all ON and OFF cells, relative to the *LL*_*x*_ of the separable GLM.

Other ganglion cells models had an opposite-sign (or “Pull”) suppression in addition to a same-sign suppression (Fig. 5C). There was in fact every combination evident in the sNIM fits to ON and OFF cells, with a majority of OFF cells having a detectable suppressive component, while slightly more than half of ON cells did not (Fig. 5D). However, the detection of suppressive RFs typically lead to much smaller improvements in model performance (Fig. 5E; median = 10.0%, *N*=290), because the effect of suppressive RFs only had a small impact on the predicted firing rate (Fig. 5B), related to its precision (Butts et al., 2007; 2011). Nevertheless, the detection of the diverse array of suppressive RFs demonstrates the ability to more accurately reflect the diversity of computation of different RF types.

Suppressive RFs were also detected by the sNIM in ON-OFF neurons. Indeed, for the example neuron described by an ON and OFF excitatory RF (Fig. 6A), a delayed suppressive RF could be added to each excitatory RF (Fig. 6B). This pattern held in a majority of ON-OFF cells (Fig. 6C), although – like with the ON and OFF cells considered earlier, the addition of suppressive RFs only led to small improvements in model performance over the already large improvements associated with the ON-OFF models without suppression (Fig. 6D; median = 14.0%). However, when the sNIM is considered as a whole, it does result in considerable model improvements relative to the separable GLM (Fig. 6E; median = 89.7%, *N*=141). Thus, the diversity of suppression is another element of the sNIM that contributes to the computational diversity of ganglion cells.

**Figure 6:**
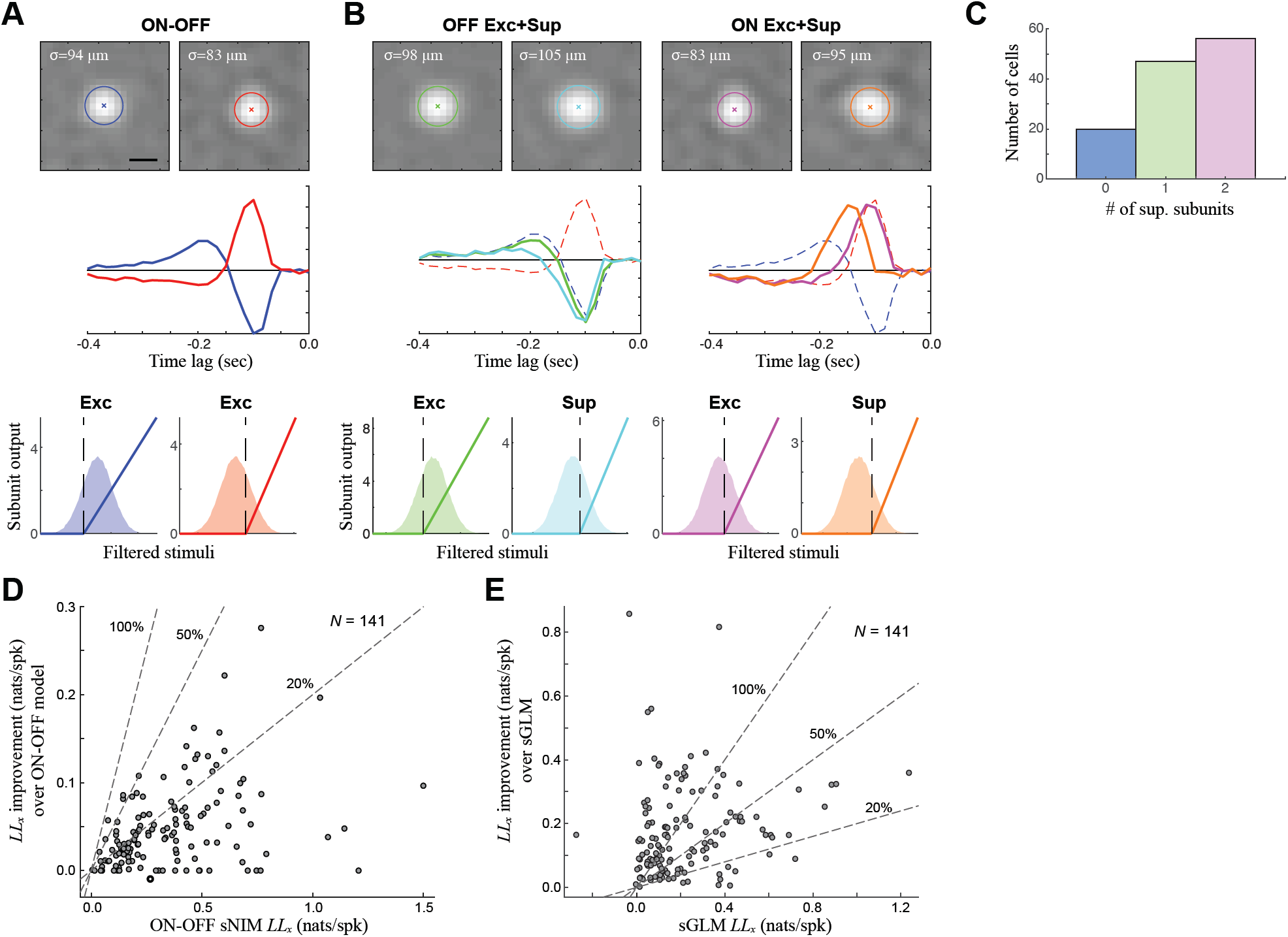
ON-OFF cells with delayed suppressive subunits. **A.** A typical ON-OFF model with two excitatory subunits can also be better modeled by the addition of suppressive subunits. **B.** A four-subunit model of the same ON-OFF cell, demonstrating a similar pattern of delayed suppression as the ON and OFF cells (Fig. 5): with each excitatory subunit having a paired, delayed, suppressive subunit. **C.** A majority of ON-OFF cells also had detectable suppressive subunits. **D.** The cross-validated log-likelihood (*LL*_*x*_) improvement of adding suppression across ON-OFF cells, relative to the *LL*_*x*_ of the ON-OFF sNIM considered in Figs. 3,4. **E.** The *LL*_*x*_-improvement of the full sNIM relative to the sGLM across ON-OFF cells.

## DISCUSSION

Here, we present a state-of-the-art modeling approach for characterizing retinal computation. It is based on extracellular recording of ganglion cell action potentials evoked by “cloud” stimuli, in which “clouds” are correlated noise (Niell and Stryker, 2008; Leon et al., 2012). Analysis of responses to clouds reveals selectivity of ganglion cells across spatial scales: here, we used a maximum likelihood estimation applied to an LNLN cascade, which is selective to multiple spatiotemporal “features” (i.e., receptive fields), to describe stimulus-driven spike train data. To handle the large number of spatial and temporal dimensions of ganglion cell RFs, we adapted fitting procedures of the Nonlinear Input Model (McFarland et al., 2013) to use space-time separable RFs (i.e. the separable NIM; sNIM). Using this experimental and computational approach, we identified a diverse array of computations in the observed ganglion cell population, which included a large number of ON-OFF cells. Our approach to characterizing ganglion cell receptive fields (RFs) can be extended to detail differences between individual retinal ganglion cell types as well as between the retinas of different species (primate, mouse, salamander, etc.).

### Comparison to previous statistical models of retinal ganglion cells

The sNIM utilizes a range of methods from existing statistical modeling methods; combining them to generate a robust description of retinal computation is novel and unique. The sNIM is based on the methods of the NIM (Butts et al., 2011; McFarland et al., 2013; Butts et al., 2016), modified to have space-time separable receptive fields (Cai et al., 1997; Butts et al., 2011; McFarland et al., 2013; Park and Pillow, 2013; Thorson et al., 2015; Maheswaranathan et al., 2018) to make it feasible to fit multiple spatiotemporal RFs per neuron at high spatial and temporal resolution. Furthermore, we employ cloud stimuli (Niell and Stryker, 2008; Leon et al., 2012) – essentially noise stimuli filtered to have spatial correlations – to drive responses to features across spatial scales. This facilitated robust fitting of salient features of ganglion cell computation, including separate ON and OFF and suppressive (i.e. inhibitory) receptive fields.

Statistical models of ganglion cells (and related LGN neurons) are some of the most successful in neuroscience, in large part due to their predominantly linear behavior (Shapley, 2009). As a result, models of ganglion cells have been largely based on spike-triggered averages (STAs) (Chichilnisky, 2001), which – although not explicitly “fit” to maximize likelihood – are the optimal linear filter in the context of stimuli such as Gaussian white noise (Chichilnisky, 2001; Dayan and Abbott, 2001). Such models, however, capture only linear stimulus processing (i.e., a single RF) and must be used with uncorrelated stimuli. Spike-triggered covariance analysis (Fairhall et al., 2006; Samengo and Gollisch, 2013; Liu and Gollisch, 2015) extended spike-triggered techniques to nonlinear models, although are only able to yield a “feature space” (e.g., Fig 2), within which the true features that the neuron is driven by exist. Thus, while such approaches identify signatures multiple excitatory and suppressive features, they cannot be used to characterize these separate components contributing to ganglion cell processing (Cantrell et al., 2010; Butts et al., 2011; McFarland et al., 2013; Liu and Gollisch, 2015).

The use of rectified nonlinearities in the context of an LNLN cascade, however, more clearly distinguishes the separate features that contributing to ganglion cell computations, due to the two-stages of nonlinear processing in such cascades better approximating the form of nonlinear processing in the retinal circuit (McFarland et al., 2013; Butts et al., 2016; Maheswaranathan et al., 2018). Such an approach was recently used in the salamander retina, where it identified multiple excitatory features comprising single OFF-receptive fields, likely corresponding to separate bipolar inputs (Maheswaranathan et al., 2018). This detection of with another recent study in salamander able to picture the spatial arrangement of such putative bipolar cell inputs in two-dimensions (Liu et al., 2017). However, neither study detected ON-OFF cells nor suppressive RFs, likely reflecting the dominant nonlinear structure of salamander RFs, which are predominantly OFF cells (Segev et al., 2006) with nonlinear excitatory integration (Gollisch and Meister, 2008a; 2008b), compared with that in the mouse retina. However, it is intriguing to believe that with more data, or tailored one-dimensional (Maheswaranathan et al., 2018) or two-dimensional (Freeman et al., 2015) stimuli, that such models could detect further dissect the distinct inputs integrated by different types of mammalian ganglion cells.

Naturally, the LNLN cascade only approximates the true nonlinearities present in ganglion cell computation, and more detailed studies have been able to model particular elements of ganglion cells that cannot be captured in this general framework. For example, early studies of the retina (Shapley and Victor, 1981) and LGN (Bonin et al., 2005; Mante et al., 2008) fit models that explicitly incorporate “contrast gain” to explain effects of contrast adaptation, which has been addressed in more recent studies using a model for synaptic depression (Jarsky et al., 2011; Ozuysal and Baccus, 2012) and presynaptic inhibition (Cui et al., 2016b). Similarly, models tailored to explain other nonlinear response properties such as temporal precision (Berry and Meister, 1998; Keat et al., 2001; Butts et al., 2011) and nonlinear spatial integration (Schwartz et al., 2012; Gollisch, 2013; Freeman et al., 2015; Liu et al., 2017) have been fit to data from limited experimental paradigms.

Detailed biophysical models taking into account the true circuitry and biophysical details of the ganglion cell circuits cannot be constrained adequately experimentally, nor can their many parameters be fit directly with data. Therefore, such phenomenological models are useful to probe more mechanistic – and yet still approximate – hypotheses about the underlying circuitry. More detailed phenomenological models – such as those considering nonlinear spatial integration (Takeshita and Gollisch, 2014; Freeman et al., 2015) can be fit in only very limited circumstances owing to the large number of component parameters and thus are not yet general. New machine-learning methods can fit large populations of recorded neurons simultaneously (McIntosh et al., 2016) and provide an opportunity to fit large numbers of parameters with more data. Such techniques, however, are still in their infancy and currently have limited interpretability. In this context, the sNIM is relatively simple: in a robust and user-friendly framework, it is able to approximate many of the nonlinearities revealed by more detailed studies. Assumptions about the specific structure of the model, such as space-time separability of the RFs, and the limit in the number of RF subunits, are necessarily approximations, but they maximize robustness and interpretability. Thus, the sNIM serves as a much-improved baseline model relative to the LN analysis that has dominated retinal computational analysis until recently.

### ON-OFF selectivity and the diversity of ganglion cell computation

While making up only a small part of ganglion cell selectivity in cat and primate (Perry et al., 1984; Troy and Shou, 2002), a significant fraction of ganglion cells in mouse are ON-OFF cells (Sun et al., 2002; Coombs et al., 2007; Zhang et al., 2012). ON-OFF cells appear to be common in salamander retina as well (Hensley et al., 1993; Gollisch and Meister, 2008b). Furthermore, an intriguing recent study utilizing LN analysis suggested that the response polarity of the linear receptive fields of mouse and pig retinal ganglion cells varied with illumination intensity (Tikidji-Hamburyan et al., 2015): simple ON or OFF characterizations of retinal neurons therefore may be insufficient to capture their physiological functionality. Similar “polarity reversal” was observed in salamander OFF RGCs in response to “peripheral image shift” (Geffen et al., 2007). This suggests that robust methods to characterize ON-OFF responses are necessary (Cantrell et al., 2010). The results we presented here demonstrates that the sNIM detects ON-OFF selectivity more sensitively than physiological characterizations using flashed stimuli and/or based on the linear RF (e.g., (Chichilnisky and Kalmar, 2002; Carcieri et al., 2003; Segev et al., 2006)). The clear RFs detected also permit detailed characterization of the measured RFs (Fig. 1) and thus provide a framework for understanding the contribution of ON-OFF processing in vision.

More broadly, the revolution in mouse genetics has allowed for the discovery of new ganglion cells – e.g., ipRGCs (Hattar et al., 2002), J-RGCs (Kim et al., 2008), and W3 cells (Zhang et al., 2012) – and the study of their presynaptic circuitry. Further, new anatomical and statistical techniques (Helmstaedter et al., 2013; Sümbül et al., 2014; Baden et al., 2016) have amplified our understanding of novel and known ganglion cell types. Nonlinear modeling approaches have offered a picture of ganglion cell diversity in more general stimulus contexts. The sNIM can be applied to general stimuli and offers a robust description of ganglion cell computation that can be related to each ganglion cell’s stimulus selectivity, a diversity of nonlinear response properties, and potentially underlying mechanisms of processing within the visual circuitry.

## METHODS

### Neurophysiology

Experimental use of mice was performed under an animal protocol approved by the IACUC of the University of Maryland. C57bl/6 mice of either sex were dark-adapted for 1-2 hrs before isoflurane anesthesia followed by euthanasia by decapitation. Eyes were removed and retinas dissected free in Ames’ medium (Sigma) bubbled with 95% O_2_/ 5% CO_2_ (Carbogen) at room temperature. Ventral retinae were isolated in small, rectangular sections and placed ganglion cells down on a 6×10 perforated multielectrode array (MEA, *ALA Scientific Instruments, Inc*). After a rest period of at least 30 minutes (to permit tissue adhesion to the MEA), ganglion cell responses to light stimuli were recorded while the tissue was perfused with Ames’ medium bubbled with Carbogen and kept at 32°C. All dissection procedures were performed under dim red illumination.

### Stimuli

Visual stimuli were generated using the *Psychophysics Toolbox* in *Matlab* and presented in the UV spectrum (*I*_mean_ ≈ 5 x 10^3^ photons/cm^2^/s, 398 nm) using a modified DLP projector (*HP Notebook Projection Companion Projector*, Model: HSTNN-FP01) (frame rate = 60 Hz), the output of which was focused through an inverted microscope objective. Stimuli include: (1) 120-s, square-wave whole-field flash at 1 Hz; (2) Gaussian white noise (GWN) flickering checkerboard (pixel size = 44.77 µm); (3) spatially correlated “cloud” stimulus; and (4) repeated, short periods of cloud (short repeats) for cross validation. The cloud stimulus was generated by low-pass filtering the GWN. The total duration of GWN, cloud, and short repeats was 20 min, and was broken into two 10-min blocks, separated by 10-min resting intervals. A 60-s long conditioning, whole-field gray light at *I*_mean_ was presented at the beginning of each stimulus (block). Data acquisition was performed using the MC_Rack software (*ALA Scientific Instruments, Inc.*), at a 50-kHz sampling rate. Spikes were sorted using their supervised offline spike sorting.

### Data Analysis

A majority of experiments and neurons were recorded using just cloud stimulation (24 preparations, 431 well-isolated ganglion cells). Due to the instability of long recordings over the two 10-min blocks of cloud stimulus and repeats, we only used both blocks to fit models for 12 of the 24 experiments, using two criteria: (1) average firing rates of neurons did not change more than 50% from the first to second experiment; (2) cross-validated likelihood of the LN model did not change by more than 50%. If either condition was not met, we used the block for which contained (on average) the best fits without suppressive terms added (LN or ON-OFF fits). In most experiments, the repeated stimulation was not stable by these criteria. However, we identified a number of individual repeat blocks for which cross-validated log-likelihoods and firing rates were sufficiently stable, but did not use these numbers in population statistics.

For a smaller set of experiments (6 experiments, 125 well-isolated neurons), we recorded neurons in both cloud and GWN conditions in order to facilitate comparisons of spatial RFs between conditions. Here, we used neurons based on having clear temporal kernels in the sGLM fits for both cloud and GWN conditions, and that the RFs did not overlap with the edge of stimulation, leaving 81 neurons. Note that these comparisons were based on comparing ON an OFF RFs (without regard to later analyses identifying ON-OFF neurons), and thus would implicitly eliminate some ON-OFF cells with strange linear RFs. Also, because performance comparisons were not important for this analysis, we did not use the stringent stability above for rejecting segments of the data.

Data was analyzed using bins at exactly twice the stimulus refresh frequency (roughly 16 ms) for all subsequent analyses.

### Linear and Nonlinear modeling

This work follows general modeling for the Nonlinear Input Model (NIM) described in detail in McFarland et al (2013); modified to incorporate separable spatiotemporal RFs. As described below, the model fitting code (in *Matlab*) has been made publicly available, and should complement the descriptions here and in previous publications. Briefly, all models were fit to maximize the penalized Poisson log-likelihood of the model given the data (Paninski, 2004), given by:

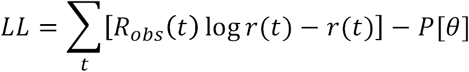

where *r*(*t*) is the predicted spike output of the model, *R*_*obs*_(*t*) is the binned spike count of the neuron being modeled, and *P*[*θ*] is a regularization penalty that depends on the model parameters *θ*.

The predicted firing rate of the GLM model is given, familiarly, by *r*(*t*) = *F*[**K**·**s**(*t*)] (Paninski, 2004; Truccolo et al., 2005) (also eq. 1, below), where we are using vector notation [**bold**] to denote the linear projection of the stimulus at a given time point (which includes all spatial positions and time lags) and the spatiotemporal filter **K**= *K*(*x,y*,**τ**) over all spatial positions and lags (back in time). The spiking nonlinearity *F*[.] is defined to be *F*[*x*]=log[1+exp(*x*)] (Paninski, 2004; McFarland et al., 2013). For the separable GLM, we approximated the full spatiotemporal kernel *K*(*x,y*,τ) as the outer product of a spatial kernel *k*_*sp*_(*x,y*) and temporal kernel *k*_*t*_(τ), resulting in the separable GLM (sGLM) used below.

The predicted firing rate of the NIM is given by (McFarland et al., 2013) (eq. 2, below). As with the sGLM, the separable NIM (sNIM) involved approximating each subunit RF as the outer product of a spatial and a temporal kernel.

Because the outer product does not uniquely specify either kernel (because one can multiply one by any constant and divide the other to get a mathematically equivalent result), we constrained the spatial kernels to be normalized (i.e., fixed overall magnitude), and to have positive centers.

Models were fit using gradient ascent of the *LL* (above), as described in detail previously (McFarland et al., 2013). However, due to the separable kernels, each RF was fit by alternating fits of the spatial component (holding the temporal component fixed) and temporal component (holding the spatial component fixed) until convergence. For each neuron, we first fit computed the spike triggered average, and then used singular value decomposition to initialize an sGLM. Once the sGLM was fit, we initialized an sNIM with two excitatory kernels, where the first was initialized from the sGLM fit, and the second had a temporal kernel multiplied by −1 (i.e., an “ON-OFF” fit). If the result had a better *LL* than the LN fit and maintained an ON-OFF form, we then fit models with additional suppressive terms. Otherwise, we generated an NIM with a single excitatory subunit (initialized using the sGLM) and added suppressive terms.

All models were regularized using a Laplacian square penalty separately on the spatial and temporal kernels – which penalizes the second derivative (McFarland et al., 2013), resulting the penalized log-likelihood (above).

### Spatial RF analysis

The robust identification of multiple spatiotemporal RFs contributing to a given ganglion cell response allows for detailed analysis of their spatial integration properties, as well as detecting the presence of surrounds. To this end, we fit circular Gaussians to a given spatial RF, fitting the precise center, standard deviation, and amplitude (4-parameters) by minimizing the mean-squared error using simplex optimization. We also could robustly fit elliptical Gaussians (two additional parameters) but found that this did not reveal additional useful information. With the centers identified, we could then take the average magnitude of pixels at any given radius from the center, generating a radial profile *w*(*r*) that revealed the center and surround. We also reported the center and surround areas, given by identifying the zero-crossing of *w*(*r*) and performing a weighted integration of (2π*r*)*w*(*r*) on either side of this zero-crossing.

### Availability of modeling code

All modeling code is available at http://neurotheory.umd.edu/code.

## References

Asari H, Meister M. Divergence of visual channels in the inner retina. Nat Neurosci 15: 1581–1589, 2012.

Baccus SA, Meister M. Fast and slow contrast adaptation in retinal circuitry. Neuron 36: 909–919, 2002.

Baden T, Berens P, Franke K, Roman-Roson M, Bethge M, Euler T. The functional diversity of retinal ganglion cells in the mouse. Nature 529: 345–350, 2016.

Barlow HB, Levick WR. The mechanism of directionally selective units in rabbit’s retina. J Physiol 178: 477–504, 1965.

Berry MJ II, Meister M. Refractoriness and neural precision. J Neurosci 18: 2200–2211, 1998.

Bonin V, Mante V, Carandini M. The suppressive field of neurons in lateral geniculate nucleus. J Neurosci 25: 10844–10856, 2005.

Bonin V, Mante V, Carandini M. The statistical computation underlying contrast gain control. J Neurosci 26: 6346–6353, 2006.

Brown SP, Masland RH. Spatial scale and cellular substrate of contrast adaptation by retinal ganglion cells. Nat Neurosci 4: 44–51, 2001.

Butts DA, Cui Y, Casti ARR. Nonlinear computations shaping temporal processing of precortical vision. J Neurophysiol 116: 1344–1357, 2016.

Butts DA, Weng C, Jin JZ, Alonso J-M, Paninski L. Temporal precision in the visual pathway through the interplay of excitation and stimulus-driven suppression. J Neurosci 31: 11313–11327, 2011.

Butts DA, Weng C, Jin JZ, Yeh C-I, Lesica NA, Alonso J-M, Stanley GB. Temporal precision in the neural code and the timescales of natural vision. Nature 449: 92–95, 2007.

Cai D, DeAngelis GC, Freeman RD. Spatiotemporal receptive field organization in the lateral geniculate nucleus of cats and kittens. J Neurophysiol 78: 1045–1061, 1997.

Cantrell DR, Cang J, Troy JB, Liu X. Non-centered spike-triggered covariance analysis reveals neurotrophin-3 as a developmental regulator of receptive field properties of ON-OFF retinal ganglion cells. PLoS Comput Biol 6: e1000967, 2010.

Carcieri SM, Jacobs AL, Nirenberg SH. Classification of retinal ganglion cells: a statistical approach. J Neurophysiol 90: 1704–1713, 2003.

Chichilnisky EJ, Kalmar RS. Functional asymmetries in ON and OFF ganglion cells of primate retina. J Neurosci 22: 2737–2747, 2002.

Chichilnisky EJ. A simple white noise analysis of neuronal light responses. Network 12: 199–213, 2001.

Coombs JL, van der List D, Chalupa LM. Morphological properties of mouse retinal ganglion cells during postnatal development. J Comp Neurol 503: 803–814, 2007.

Cui Y, Liu LD, McFarland JM, Pack CC, Butts DA. Inferring Cortical Variability from Local Field Potentials. J Neurosci 36: 4121–4135, 2016a.

Cui Y, Wang YV, Park S, Demb JB, Butts DA. Divisive suppression explains high-precision firing and contrast adaptation in retinal ganglion cells. Elife (2016b). doi: 10.7554/eLife.19460.001.

Dayan P, Abbott LF. Theoretical neuroscience: computational and mathematical modeling of neural systems. Cambridge, MA: Massachusetts Institute of Technology Press, 2001.

Demb JB, Singer JH. Functional Circuitry of the Retina. Annu Rev Vis Sci 1: 263–289, 2015.

Dhande OS, Huberman AD. Retinal ganglion cell maps in the brain: implications for visual processing. Curr Opin Neurobio 24: 133–142, 2014.

Dhande OS, Stafford BK, Lim J-HA, Huberman AD. Contributions of Retinal Ganglion Cells to Subcortical Visual Processing and Behaviors. Annu Rev Vis Sci 1: 291–328, 2015.

Euler T, Haverkamp S, Schubert T. Retinal bipolar cells: elementary building blocks of vision. Nat Rev Neurosci 15: 507–519, 2014.

Fairhall AL, Burlingame CA, Narasimhan R, Harris RA, Puchalla JL, Berry MJ II. Selectivity for multiple stimulus features in retinal ganglion cells. J Neurophysiol 96: 2724–2738, 2006.

Freeman J, Field GD, Li PH, Greschner M, Gunning DE. Mapping nonlinear receptive field structure in primate retina at single cone resolution. Elife (2015). doi: 10.7554/eLife.05241.001.

Geffen MN, de Vries SEJ, Meister M. Retinal ganglion cells can rapidly change polarity from Off to On. PLoS Biol 5: e65, 2007.

Gollisch T, Meister M. Modeling convergent ON and OFF pathways in the early visual system. Biological Cybernetics 99: 263–278, 2008a.

Gollisch T, Meister M. Rapid neural coding in the retina with relative spike latencies. Science 319: 1108–1111, 2008b.

Gollisch T. Features and functions of nonlinear spatial integration by retinal ganglion cells. J Physiol-Paris 107: 338–348, 2013.

Hattar S, Liao HW, Takao M, Berson DM, Yau KW. Melanopsin-containing retinal ganglion cells: architecture, projections, and intrinsic photosensitivity. Science 295: 1065–1070, 2002.

Helmstaedter M, Briggman KL, Turaga SC, Jain V, Seung HS, Denk W. Connectomic reconstruction of the inner plexiform layer in the mouse retina. Nature 500: 168–174, 2013.

Hensley SH, Yang XL, Wu SM. Relative contribution of rod and cone inputs to bipolar cells and ganglion cells in the tiger salamander retina. J Neurophysiol 69: 2086–2098, 1993.

Hochstein S, Shapley RM. Linear and nonlinear spatial subunits in Y cat retinal ganglion cells. J Physiol 262: 265–284, 1976.

Jarsky T, Cembrowski MS, Logan SM, Kath WL, Riecke H, Demb JB, Singer JH. A synaptic mechanism for retinal adaptation to luminance and contrast. J Neurosci 31: 11003–11015, 2011.

Keat J, Reinagel P, Reid RC, Meister M. Predicting every spike: a model for the responses of visual neurons. Neuron 30: 803–817, 2001.

Kim DS, Ross SE, Trimarchi JM, Aach J, Greenberg ME, Cepko CL. Identification of molecular markers of bipolar cells in the murine retina. J Comp Neurol 507: 1795–1810, 2008.

Kim KJ, Rieke F. Temporal contrast adaptation in the input and output signals of salamander retinal ganglion cells. J Neurosci 21: 287–299, 2001.

Korenberg MJ, Sakai HM, Naka K-I. Dissection of the neuron network in the catfish inner retina. III. Interpretation of spike kernels. J Neurophysiol 61: 1110–1120, 1989.

Leon PS, Vanzetta I, Masson GS, Perrinet LU. Motion clouds: model-based stimulus synthesis of natural-like random textures for the study of motion perception. J Neurophysiol 107: 3217–3226, 2012.

Lettvin JY, Maturana HR, the WMPO, 1959. What the frog“s eye tells the frog”s brain. Proc IEEE 47: 1940–1951, 1959.

Liu JK, Gollisch T. Spike-Triggered Covariance Analysis Reveals Phenomenological Diversity of Contrast Adaptation in the Retina. PLoS Comput Biol 11: e1004425, 2015.

Liu JK, Schreyer HM, Onken A, Rozenblit F, Khani MH, Krishnamoorthy V, Panzeri S, Gollisch T. Inference of neuronal functional circuitry with spike-triggered non-negative matrix factorization. Nat Commun 8: 149, 2017.

Liu RC, Tzonev S, Rebrik SP, Miller KD. Variability and information in a neural code of the cat lateral geniculate nucleus. J Neurophysiol 86: 2789–2806, 2001.

Maheswaranathan N, Kastner DB, Baccus SA, Ganguli S. Inferring hidden structure in multilayered neural circuits. PLoS Comput Biol 14: e1006291, 2018.

Mante V, Bonin V, Carandini M. Functional mechanisms shaping lateral geniculate responses to artificial and natural stimuli. Neuron 58: 625–638, 2008.

Masland RH. The neuronal organization of the retina. Neuron 76: 266–280, 2012.

McFarland JM, Cui Y, Butts DA. Inferring nonlinear neuronal computation based on physiologically plausible inputs. PLoS Comput Biol 9: e1003143, 2013.

McIntosh LT, Maheswaranathan N, Nayebi A, Ganguli S, Baccus SA. Deep Learning Models of the Retinal Response to Natural Scenes. Adv Neural Inf Process Syst 29: 1369–1377, 2016.

Meister M, Pine J, Baylor DA. Multi-neuronal signals from the retina: acquisition and analysis. J Neurosci Methods 51: 95–106, 1994.

Münch TA, da Silveira RA, Siegert S, Viney TJ, Awatramani GB, Roska B. Approach sensitivity in the retina processed by a multifunctional neural circuit. Nat Neurosci 12: 1308–1316, 2009.

Niell CM, Stryker MP. Highly selective receptive fields in mouse visual cortex. J Neurosci 28: 7520–7536, 2008.

Olveczky BP, Baccus SA, Meister M. Segregation of object and background motion in the retina. Nature 423: 401–408, 2003.

Ozuysal Y, Baccus SA. Linking the computational structure of variance adaptation to biophysical mechanisms. Neuron 73: 1002–1015, 2012.

Pamplona D, Hilgen G, Cessac B, Sernagor E, Kornprobst P. A super-resolution approach for receptive fields estimation of neuronal ensembles. BMC Neurosci 16: P130, 2015.

Paninski L. Maximum likelihood estimation of cascade point-process neural encoding models. Network 15: 243–262, 2004.

Park M, Pillow JW. Bayesian inference for low rank spatiotemporal neural receptive fields. Advances in Neural Information Processing ….

Perry VH, Oehler R, Cowey A. Retinal ganglion cells that project to the dorsal lateral geniculate nucleus in the macaque monkey. Neuroscience 12: 1101–1123, 1984.

Pillow JW, Paninski L, Uzzell VJ, Simoncelli EP, Chichilnisky EJ. Prediction and decoding of retinal ganglion cell responses with a probabilistic spiking model. J Neurosci 25: 11003–11013, 2005.

Samengo I, Gollisch T. Spike-triggered covariance: geometric proof, symmetry properties, and extension beyond Gaussian stimuli. J Comput Neurosci 34: 137–161, 2013.

Schwartz GW, Okawa H, Dunn FA, Morgan JL, Kerschensteiner D, Wong ROL, Rieke F. The spatial structure of a nonlinear receptive field. Nat Neurosci 15: 1572–1580, 2012.

Segev R, Puchalla JL, Berry MJ II. Functional organization of ganglion cells in the salamander retina. J Neurophysiol 95: 2277–2292, 2006.

Seung HS, Sümbül U. Neuronal cell types and connectivity: lessons from the retina. Neuron 83: 1262–1272, 2014.

Shapley RM, Victor JD. The effect of contrast on the transfer properties of cat retinal ganglion cells. J Physiol 285: 275–298, 1978.

Shapley RM, Victor JD. How the contrast gain control modifies the frequency responses of cat retinal ganglion cells. J Physiol 318: 161–179, 1981.

Shapley RM. Linear and nonlinear systems analysis of the visual system: why does it seem so linear? A review dedicated to the memory of Henk Spekreijse. Vision Research 49: 907–921, 2009.

Sun W, Li N, He S. Large-scale morphological survey of mouse retinal ganglion cells. J Comp Neurol 451: 115–126, 2002.

Sümbül U, Song S, McCulloch K, Becker M, Lin B, Sanes JR, Masland RH, Seung HS. A genetic and computational approach to structurally classify neuronal types. Nat Commun 5: 3512, 2014.

Takeshita D, Gollisch T. Nonlinear spatial integration in the receptive field surround of retinal ganglion cells. J Neurosci 34: 7548–7561, 2014.

Theunissen FE, David SV, Singh NC, Hsu A, Vinje WE, Gallant JL. Estimating spatio-temporal receptive fields of auditory and visual neurons from their responses to natural stimuli. Network 12: 289–316, 2001.

Thorson IL, Liénard J, David SV. The Essential Complexity of Auditory Receptive Fields. PLoS Comput Biol 11: e1004628, 2015.

Tikidji-Hamburyan A, Reinhard K, Seitter H, Hovhannisyan A, Procyk CA, Allen AE, Schenk M, Lucas RJ, Münch TA. Retinal output changes qualitatively with every change in ambient illuminance. Nat Neurosci 18: 66–74, 2015.

Troy JB, Shou T. The receptive fields of cat retinal ganglion cells in physiological and pathological states: where we are after half a century of research. Prog Retin Eye Res 21: 263–302, 2002.

Truccolo W, Eden UT, Fellows MR, Donoghue JP, Brown EN. A point process framework for relating neural spiking activity to spiking history, neural ensemble, and extrinsic covariate effects. J Neurophysiol 93: 1074–1089, 2005.

Uzzell VJ, Chichilnisky EJ. Precision of spike trains in primate retinal ganglion cells. J Neurophysiol 92: 780–789, 2004.

Zaghloul KA, Boahen K, Demb JB. Contrast adaptation in subthreshold and spiking responses of mammalian Y-type retinal ganglion cells. J Neurosci 25: 860–868, 2005.

Zhang Y, Kim I-J, Sanes JR, Meister M. The most numerous ganglion cell type of the mouse retina is a selective feature detector. PNAS 109: E2391–8, 2012.

